# Embryo-specific epigenetic mechanisms reconstitute the CHH methylation landscape during Arabidopsis embryogenesis

**DOI:** 10.1101/2023.04.06.535361

**Authors:** Ping-Hung Hsieh, Jennifer M. Frost, Yeonhee Choi, Tzung-Fu Hsieh, Daniel Zilberman, Robert L Fischer

## Abstract

The modification of flowering plant DNA by CHH methylation acts primarily to silence transposable elements, of which many active copies are present in *Arabidopsis thaliana*. During embryogenesis, the CHH methylation landscape is dramatically reprogrammed, resulting in exceedingly high levels of this modification upon mature embryo formation. The mechanisms constituting the remodeling process, and its function in embryos, are unclear. Here, we isolate embryos from Arabidopsis plants harboring mutations for key components of the pathways that confer CHH methylation, namely RNA-directed DNA methylation (RdDM) and the Chromomethylase 2 (CMT2) pathways. We reveal that embryos are more methylated than leaves at shared CMT2 and RdDM targeting loci, accounting for most embryonic CHH hypermethylation. While the majority of embryo CHH methylated loci overlap with those in somatic tissues, a subset of conventional pericentric CMT2-methylated loci are instead targeted by RdDM in embryos. These loci, termed ‘embRdDM’ exhibit intermediate H3K9me2 levels, associated with increased chromatin accessibility. Strikingly, more than 50% of the embRdDM loci in pollen vegetative (nurse) cells and *ddm1* mutant somatic tissues are also targeted by RdDM, and these tissues were also reported to exhibit increased chromatin accessibility in pericentric heterochromatin. Furthermore, the root columella stem cell niche also displays CHH hypermethylation and an enriched presence of small RNAs at embRdDM loci. Finally, we observe a significant overlap of CHH hypermethylated loci with endosperm DEMETER targeting sites, suggesting that non-cell autonomous communication within the seed may contribute to the epigenetic landscape of the embryo. However, similar overlap with vegetative cell DEMETER targets indicates that the chromatin landscape that allows DEMETER access is mirrored in developing embryos, permitting CHH methylation catalysis at the same loci. Our findings demonstrate that both conserved and embryo-specific epigenetic mechanisms reshape CHH methylation profiles in the dynamic chromatin environment of embryogenesis.

## Introduction

In flowering plants, the majority of DNA methylation is located at transposable elements (TEs). TEs make up a variable proportion of plant genomes, and Arabidopsis is relatively TE poor, but retains several families of active elements that must be repressed to prevent potentially deleterious transposition events (Tsukahara et al. 2009). In general, TE DNA in plants is enriched with non-CG methylation, catalysed by two pathways. The RNA-directed DNA methylation (RdDM) pathway predominates at TE edges and in pericentric chromatin, with low levels of histone H3K9 dimethylation (H3K9me2). In brief, RdDM is initiated by Pol IV transcription of single-stranded RNAs, utilised by RNA-DEPENDENT RNA POLYMERASE 2 (RDR2) to produce double stranded RNAs. Pol IV association with DNA is mediated by chromatin remodellers CLASSY1-4 (CLSY1-4) which confer locus- and tissue-specificity (Zhou et al. 2018). Double-stranded RNAs (dsRNAs) are then processed by DICER-LIKE 3 (DCL3) into 24-nucleotide siRNAs and incorporated onto ARGONAUTE 4 (AGO4), which recruits DOMAINS REARRANGED METHYLTRANSFERASE 2 (DRM2) to the locus, catalysing DNA methylation. In TE bodies, and heterochromatic regions with high H3K9me2, methylation is mediated by CHROMOMETHYLASE2 (CMT2) and CMT3 at CHH and CHG contexts respectively (Law and Jacobsen 2010; Hsieh et al. 2016; Stroud et al. 2014).

TE silencing is especially important in tissues that contribute to future generations, including gametes, stem cell niches, such as the shoot and root apical meristems, and the developing embryo. Comprehensive mechanisms of TE silencing are found in these tissues, almost all based on RdDM (Kawakatsu et al. 2016; Baubec et al. 2014). For example, in the male germline, siRNAs generated by tapetum cells move into meiocytes, initiating silencing of TEs via RdDM (Long et al. 2021). Then, during gametogenesis, gamete companion cells undergo deep DNA CG demethylation by the DEMETER glycosylase (Hsieh et al. 2009; Ibarra et al. 2012; Schoft et al. 2011; Park et al. 2016). This results in further non-cell autonomous RdDM activity, at least in pollen, where siRNAs generated in the vegetative cell lead to TE repression in sperm (Slotkin et al. 2009; Martínez et al. 2016).

In the embryo, CHH methylation builds up during development in both euchromatic and heterochromatic TEs, which leads to exceptionally high levels of CHH methylation that peak at maturation, with levels being much lower post germination (Bouyer et al. 2017; Papareddy et al. 2020; Kawakatsu et al. 2017). The embryo develops within the seed, in close contact with the endosperm, the fertilized product of the CG demethylated central cell. During seed development, the endosperm generates abundant siRNAs (Mosher et al. 2009) leading to speculation that endosperm siRNAs may move into the embryo and reinforce TE silencing, in a synonymous mechanism to that of pollen. Indeed, CHH hypermethylation in embryo is linked to high siRNA levels (Papareddy et al. 2020), however, the mechanisms behind high embryonic CHH methylation have not been fully elucidated. Here, we set out to characterize the molecular pathways that contribute to embryonic CHH methylation, as well as the role of this process in Arabidopsis reproduction and development.

## Results

### Mechanisms of CHH DNA methylation maintenance are similar between leaves, vegetative cells and embryos

To assess the contribution of CMT2 and RdDM pathways to the establishment of CHH hypermethylation during embryogenesis, we isolated *cmt2*, *rdr2rdr6* (*r2r6*) and wild-type (WT) *Arabidopsis* embryos at early linear cotyledon (lcot) and mature green (mg) stages and analyzed their methylomes by whole-genome bisulfite-sequencing. Using published sperm methylome data (Hsieh et al., 2016) as a proxy of the initial CHH methylation profile pre-embryogenesis, we found that in *cmt2* mutant embryos the increase in CHH methylation, as conferred by RdDM, was far more profound from early lcot to mature embryo development than that observed in the transition from sperm to early lcot embryo (Fig.1A). This result suggests that RdDM-mediated CHH methylation mainly occurs during the early lcot-to-mature embryogenesis period. Conversely, in *r2r6* embryos, CMT2-mediated CHH methylation was found to occur during fertilization and continue through embryogenesis (Fig.1B). The discrete patterns of CHH methylation increase between the two pathways thus suggest that most CMT2 targets become methylated during morphogenesis stages, while most RdDM targets are methylated during maturation phase of embryogenesis.

**Fig. 1.**
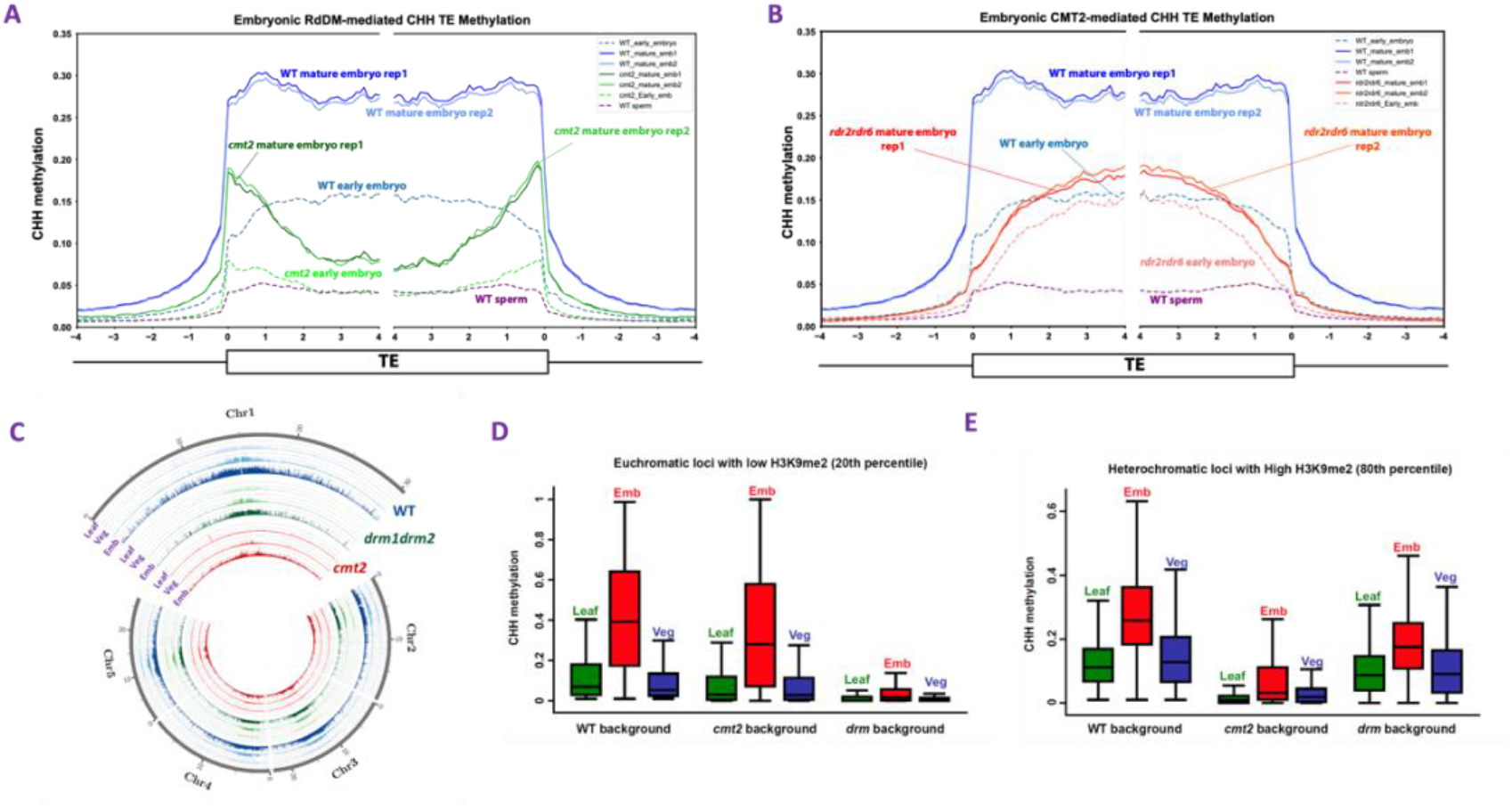
TE CHH methylation is regulated similarly in leaves, vegetative cells, and embryos. Re-establishment of TE CHH methylation in embryos via (A) RdDM-dependent and (B) CMT2-dependent pathways. Methylation within each 100-bp interval was averaged and plotted from 4 kb away from the annotated TE (negative numbers) to 4 kb into the annotated region (positive numbers). (C) Genome-wide CHH methylation levels in leaves, vegetative cells, and embryos of WT, *drm* and *cmt2* plants. CHH methylation levels were calculated in 10-kb windows across the genome. Box plots show CHH methylation levels within 50-bp windows in (D) leaf euchromatic loci with low H3K9me2 levels (H3K9me2 levels are less than the 20^th^ percentile of all windows) and (E) leaf heterochromatic loci with high H3K9me2 levels (H3K9me2 levels are greater than the 80^th^ percentile of all windows).

Next, to investigate how CHH hypermethylation is maintained in mature embryos, we analyzed CHH methylation levels of WT, *cmt2*, and *drm1drm2* (*drm*) leaves (Stroud et al. 2014; Stroud et al. 2013), and also performed methylome analysis of mature *drm1drm2* (*drm*) embryos. In the male gametophyte, the vegetative cell exhibits strikingly high levels of CHH methylation (Ibarra et al., 2012). To determine whether CHH hypermethylation is maintained similarly in mature embryos and vegetative cells we also analyzed vegetative cells in WT, *cmt2*, and *drm* genetic backgrounds (Hsieh et al. 2016). Our meta-analysis of CHH methylation levels in TEs in leaves, vegetative cells and mature embryos showed a similar pattern across tissues, whereby the *cmt2* mutation caused substantial CHH methylation loss in TE bodies, and the *drm* mutation caused substantial loss of CHH methylation at TE edges (Fig. S1A-C). The loss of CHH methylation due to the CMT2 mutation is most profound in leaves, though the overall level of CHH methylation is lower in leaves than other tissues (Fig S1A-C). Interestingly, DRM activity was not restricted to euchromatic TE edges in mature embryos and vegetative cells, since remaining DRM-mediated CHH methylation clearly extended into heterochromatic TE bodies in *cmt2* embryos and vegetative cells (Fig. S1B and S1C). None of these mutations obviously affected embryonic CG methylation levels in TEs, as expected (Fig S2A), although we found slightly elevated CG methylation in genes in *drm* mutant embryos (Fig S2B). Loss of CMT2 and DRM also resulted in a slight decrease of embryonic CHG methylation in TEs, as found in a previous study of seedlings (Fig S2C) (Zemach et al. 2013) with little effect on gene bodies (Fig S2D)

A previous study in our lab showed that CHH hypermethylation in vegetative cells occurs mostly in heterochromatic regions and is mediated by CMT2 (Hsieh et al. 2016). At the chromosome-level in embryos, we found that both CMT2- and DRM (RdDM)-mediated CHH hypermethylated regions were more deeply methylated and extended further outside of pericentromeric heterochromatin than either leaves or vegetative cells (Fig. 1C). To assess whether both CMT2- and RdDM-mediated CHH methylation were enhanced at euchromatic and heterochromatic loci in embryos, we classified the genomic loci into ‘Euchromatic Loci’ with low H3K9me2 (below the 20th percentile of all informative 50-bp genomic windows) in leaves and ‘Heterochromatic Loci’ with high H3K9me2 (above the 80th percentile) in leaves. In all three tissues, euchromatic CHH methylation was predominately dependent on DRM (RdDM). Vegetative cells and leaves had comparable RdDM-mediated and CMT2-mediated CHH methylation levels to each other in euchromatic regions (Fig. 1D; *cmt2* and *drm* mutant backgrounds, respectively), however, mature embryos showed much higher euchromatic RdDM-mediated CHH methylation and slightly higher euchromatic CMT2-mediated CHH methylation then either leaves or vegetative cells (Fig. 1D). Heterochromatic CHH methylation was primarily dependent on CMT2 in all three tissues (Fig. 1E; *drm* mutant background), but vegetative cells had slightly higher heterochromatic RdDM-mediated CHH methylation than leaves (Fig. 1E; *cmt2* mutant background), which is consistent with our previous study (Hsieh et al. 2016). Mature embryos exhibited not only higher heterochromatic RdDM-mediated CHH methylation than leaves and vegetative cells, but also higher heterochromatic CMT2-mediated CHH methylation (Fig. 1E; *cmt2* and *drm* mutant methylomes, respectively).

### Embryonic CHH hypermethylation occurs at both newly methylated embryonic sites and existing leaf RdDM and CMT2 targeting sites

Genome-wide CHH hypermethylation in embryos compared to leaves could result from either: 1) newly methylated (or at least newly detectable as methylated) loci that are unique to mature embryos; or 2) Increased methylation at leaf methylated sites. To assess the contribution of these two scenarios to “embryonic CHH hypermethylated loci”, we first extracted loci where fractional embryo CHH methylation is at least 0.1 more than leaf CHH methylation (Fig. 2A). We then categorized these embryonic CHH hypermethylated 50-bp windows into “leaf ummethylated loci” (<1% CHH methylation in WT leaves) and “leaf methylated loci” (>=1% CHH methylation) based on the CHH methylation levels of WT leaves (Methods, Fig. 2B-D). Finally, both “leaf unmethylated loci” and “leaf methylated loci” were further classified into leaf-DRM-dominant/emb-DRM-dominant, leaf-CMT2-dominant/emb-CMT2-dominant, and leaf-DRM-CMT2 co-targeting/emb-DRM-CMT2 co-targeted loci where the loci are mainly targeted by DRM, CMT2, or DRM-CMT co-targeting mechanisms in leaves and embryos, respectively (Fig 2C&D, Methods).

**Fig. 2.**
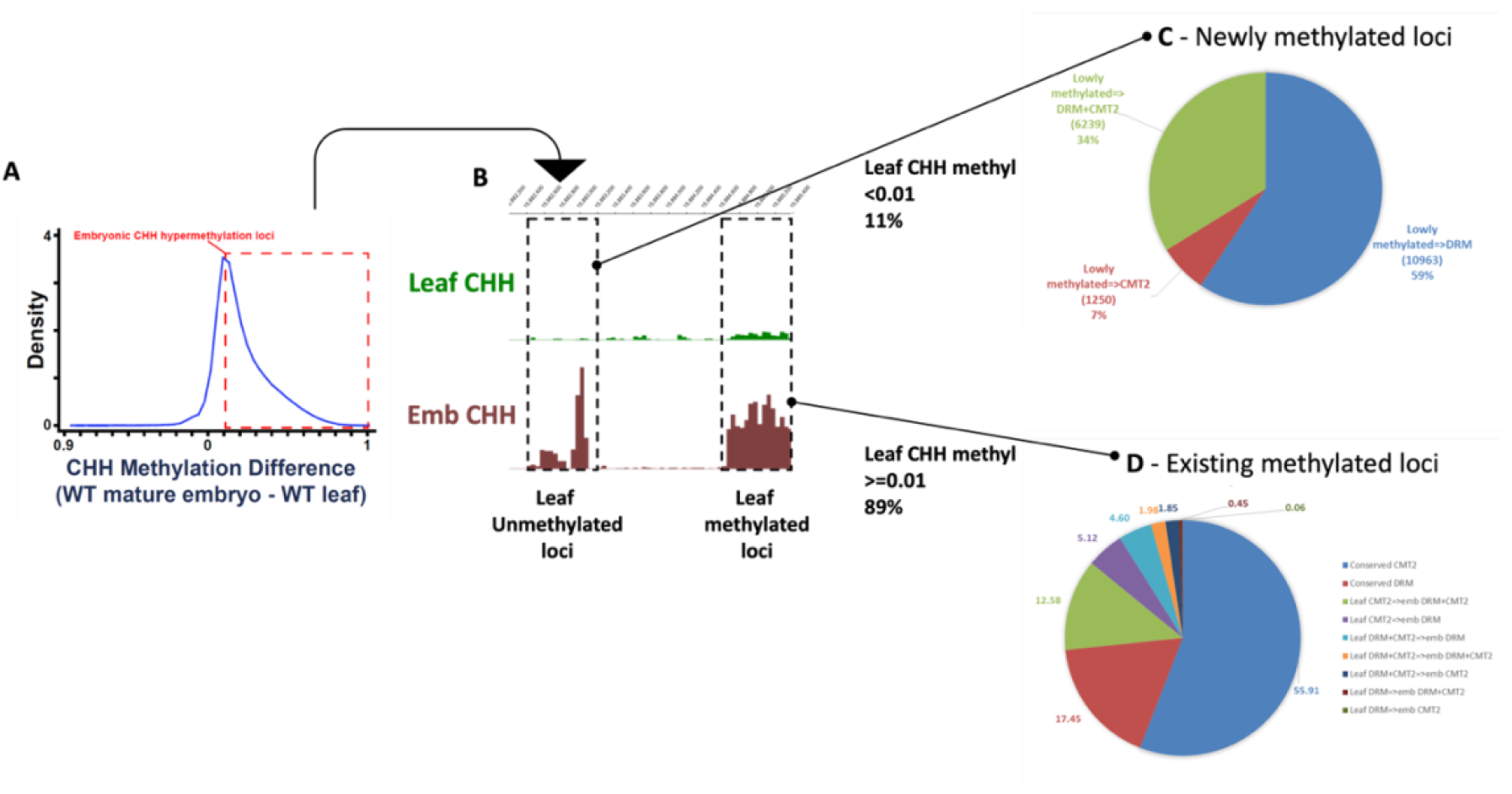
Embryonic CHH hypermethylation loci comprise both existing leaf CHH methylation targeting sites as well as new, unshared sites. (A) Kernel density plot showing the probability density of CHH DNA methylation differences between 50-bp windows retaining at least 10% CHH methylation in either mature WT embryos or leaves. Only windows with more than 20 informative sequenced cytosines in both samples are included. The dashed box indicates embryonic CHH hypermethylation loci where CHH methylation is at least 10% greater in mature WT embryos than in WT leaves. (B) Snapshot of embryonic CHH hypermethylation loci based on their methylation levels in leaves. Newly methylated (or previously undetectable) loci are where the CHH methylation levels are less than 1% in leaves. (C) Pie charts showing mechanisms that mediate embryonic CHH hypermethylation in newly methylated loci (11% of embryonic CHH hypermethylation) or (D) the remaining embryonic CHH hypermethylated loci (89 %), corresponded to existing methylated loci between leaves and embryos.

To ensure our classification of CHH methylation targeting was robust, we analyzed CHH methylation and H3K9me2 enrichment of these defined loci in mutant tissues, confirming that DRM-dominant loci lost most CHH methylation in the *drm* background, CMT2-dominant loci lost most CHH methylation in the *cmt2* background, and that *drm* and *cmt2* caused similar levels of CHH methylation loss at DRM-CMT2 co-targeted loci in leaves and embryos (Fig S3A and 3B). As expected, DRM-dominant loci had the lowest H3K9me2 levels, CMT2-dominant loci had the highest H3K9me2 levels, and DRM-CMT2 co-targeted loci had intermediate H3K9me2 levels in both leaves and embryos (Fig S3C&3D). Using this classification of CHH methylation mechanisms, we could now address the question of whether embryonic CHH hypermethylated loci were achieved by distinct mechanisms between leaves and embryos.

We found that 11% of embryonic CHH hypermethylation corresponded to “leaf unmethylated loci” (Table S1). For these novel embryonic methylated sites, *de novo* CHH methylation was predominantly catalyzed through emb-DRM-dominant and emb-DRM-CMT2 co-targeting machinery (Fig 2C, blue and green). The remaining embryonic CHH hypermethylated loci (89 %), corresponded to “leaf methylated loci”. For the majority of these loci, the same machinery conferred CHH methylation between embryos and leaves, where 55.91% were catalyzed by CMT2-targeting machinery and 17.45% by DRM-targeting machinery (Fig. 2D), indicating high levels of conservation of CHH methylation mechanisms between these tissues (Table S1). The remaining embryonic CHH hypermethylation at shared leaf methylated loci exhibited a targeting machinery switch mainly from CMT2-mediated CHH methylation in leaves to RdDM-dominant (5.12%) and RdDM-CMT2 co-targeted (12.58%) machinery in embryos (Fig. 2D, green and purple).

In summary, some embryo CHH hypermethylation is derived from a unique set of embryo-specific CHH methylated sites, which we identify as being targeted by both RdDM and RdDM-CMT2 pathways (Fig. S4). However, since the majority of CHH hypermethylated sites are still methylated in leaves, CHH methylation levels at those loci are simply higher in embryos. In most cases, the machinery contributing to embryonic hypermethylation, i.e. DRM- and CMT2-pathways, are shared between embryos and leaves, although a subset of shared loci were methylated by unique machinery in embryos, switching from CMT2-only to RdDM-CMT2 and RdDM-only mechanisms.

### Novel embryonic RdDM targets occur at loci with intermediate H3K9me2, likely due to increased chromatin accessibility during early embryogenesis

Previous studies have shown that cell type-specific chromatin environments can result in cell type-specific RdDM activities (Zhou et al. 2018; Zhou et al. 2022; Kawakatsu et al. 2016). We therefore wondered if embryo-specific RdDM targets were associated with any unique properties of embryonic chromatin. We defined “novel embryonic RdDM (embRdDM) loci” comprising (Table S1): 1) 50-bp windows that are unmethylated in leaves and are emb-DRM-dominant targets (10,963 windows) or emb-DRM-CMT2 co-targets (6,239 windows); and 2) 50-bp windows that are CMT2-dominant in leaves but become emb-DRM-dominant targets (7,798 windows) or emb-DRM-CMT2 co-targets (19,168 windows).

We found that embRdDM loci are enriched in pericentric regions (Fig. 3A), and 24.1% of leaf-CMT2-dominant sites become emb-DRM-dominant (7%, Table S1) and emb-DRM-CMT2 co-targeted loci (17.1%, Table S1). Therefore, we speculated that some of the leaf heterochromatic CMT2 targeted regions might have become accessible to RdDM in embryos. To characterize the chromatin features of embRdDM loci, we compared embryonic and leaf H3K9me2 levels (Parent et al. 2021; Stroud et al. 2014) of conserved leaf-embryo DRM loci (consvRdDM), conserved leaf-embryo CMT2 loci (consvCMT2), and embRdDM loci. We found that both embryonic and leaf H3K9me2 levels of embRdDM loci were intermediate between the low levels of consvRdDM loci and the high levels of consvCMT2 loci (Fig. 3B). The intermediate embryonic H3K9me2 levels of embRdDM loci suggest that chromatin features other than H3K9me2 enrichment are responsible for conferring the difference between leaves an embryos in RdDM accessibility. For example, the CLASSY (CLSY) chromatin remodeling factors could facilitate access of RdDM as they were found to be associated with small RNA production during embryogenesis (Papareddy et al. 2020). However, embRdDM loci were likely still not as accessible as the consvRdDM loci in embryonic cells since CHH methylation levels of embRdDM loci were lower than consvRdDM loci during embryogenesis (Fig. 3C).

**Fig. 3.**
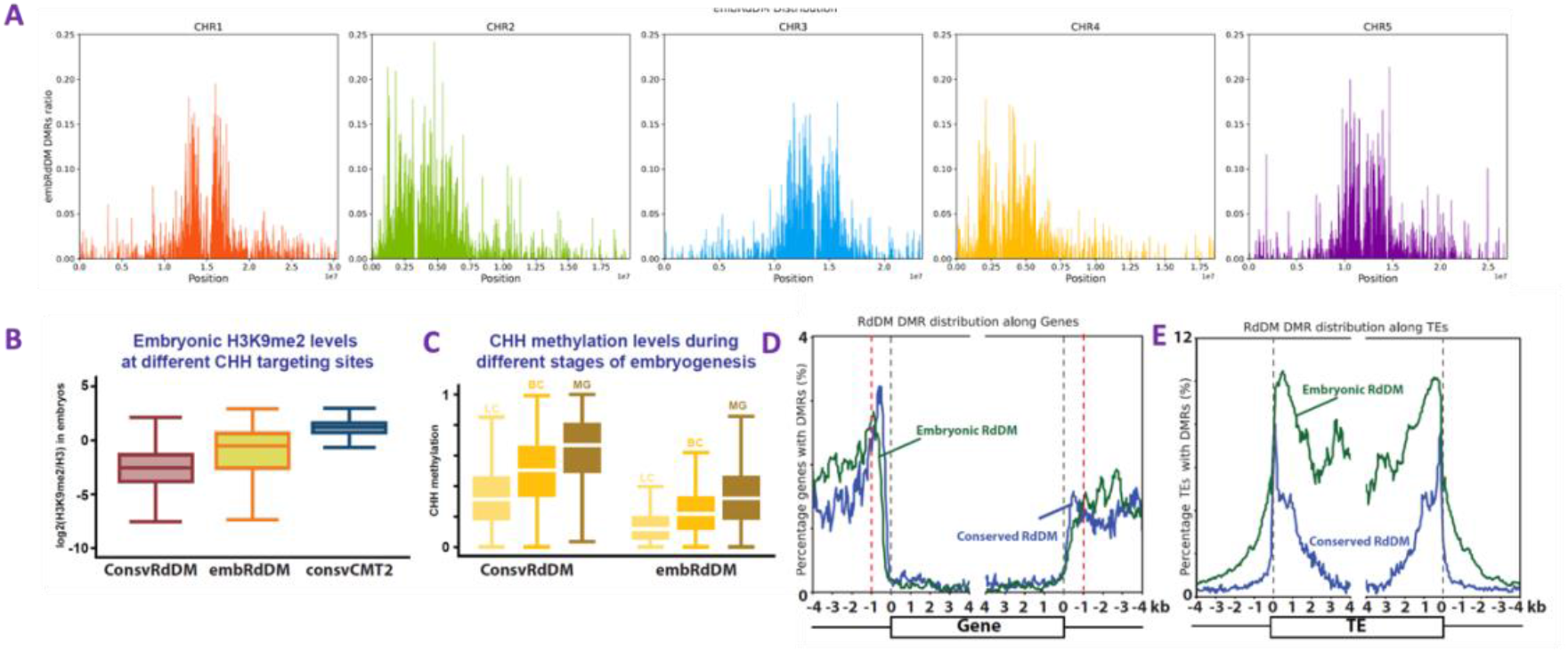
New embryo RdDM sites have medium leaf/embryo H3K9me2 levels and are RDR2-dependent. (A) Distribution of embRdDM loci along chromosomes. Y-axis values indicate the ratio of embRdDM loci in a 10kb window (B) Embryonic H3K9me2 levels in conserved RdDM (consvRdDM), embRdDM, and conserved CMT2 (consvCMT2) targeting loci. (C) CHH methylation levels in consvRdDM and embRdDM in different stages during embryogenesis (stage labels: LC = linear cotyledon, BC= bending cotyledon, MG = mature green). Distribution of embRdDM loci along (D) genes and (E) TEs. The percentage of genes and TEs with an embRdDM locus present is plotted for each 50-bp interval within 4 kb of the alignment sites. The grey dashed lines indicate transcriptional start sites and termination sites of genes. The red dashed lines indicate up- and down-stream 1 kb regions of genes.

To understand the characteristics of embRdDM loci, we next investigated whether embRdDM loci are distributed differently from consvRdDM loci in genes and TEs (Fig. 3D). ConsvRdDM loci were enriched around 500 bp upstream of genes, which was similar to embRdDM loci, though from around 1000 bp upstream embRdDM loci were more highly enriched. For TEs, consvRdDM loci preferentially targeted euchromatic TE edges, but surprisingly, embRdDM loci also showed apparent enrichment in heterochromatic TE bodies, which are usually targeted by CMT2 in leaves (Fig. 3E).

The reprogramming of CHH methylation during embryogenesis occurs concurrently with gene expression reprogramming, whereby genes can be classified in to 4 clusters: (A) highest during the pre-cotyledon phase, (B) decreasing between transition and mature phases, (C) highest during the transition phase, and (D) highest during the mature phase (Choi et al. 2021; Choi et al. 2020). To determine whether embRdDM or consvRdDM CHH methylation loci were associated with gene expression patterns, we analyzed the distribution of embRdDM and consvRdDM loci along gene clusters A-D. We found that neither consvRdDM nor embRdDM loci were associated with gene expression patterns during embryogenesis (Fig S5). However, we did find that embRdDM loci were overlapped with the transcriptional start site (TSS) of 12 genes out of 3197 genes in cluster A (highest during the pre-cotyledon phase), and speculate that TSS CHH methylation might be directly associated with the down-regulation of only these genes during embryogenesis.

Taken together, our data indicate that the incidence of embRdDM loci is likely due to a subset of CMT2-targeted genomic loci, with intermediate H3K9me2 levels in leaves, becoming accessible to RdDM in embryos during embryogenesis.

### EmbRdDM loci are RDR2-dependent in embryos and partially overlap with RdDM targets in vegetative cells and *ddm1* somatic tissues

The presence of embRdDM loci is reminiscent of the extended RdDM activity into heterochromatin that occurs in wild type vegetative cells, associated with heterochromatin decondensation and the derepression of TEs (Schoft et al. 2009; Slotkin et al. 2009). The DDM1 chromatin remodeler is required to allow access of DNA methyltransferases MET1, CMT3 and CMT2 to deposit DNA methylation in heterochromatin. *ddm1* mutations also result in extended RdDM activity into heterochromatin in somatic tissues, accompanied by heterochromatin decondensation and the derepression of TEs (Zemach et al. 2013). However, it is not known whether these distinct occurrences of CHH hypermethylation share maintenance or targeting mechanisms. In order to assess the degree of overlap between new RdDM targeted loci in embryos, vegetative cells and *ddm1* mutant somatic tissues, we compared the RdDM-mediated CHH methylation present in wild-type, *cmt2* and *ddm1* mutant tissues. As expected, embryonic CHH methylation is mainly affected by RdDM (i.e. *rdr2* mutant embryos, Fig. 4A). We observed that CHH methylation of embRdDM loci is mostly maintained by CMT2 (i.e. CMT2-dominant) in WT flowers and shoots (Fig. 4B and 4C). However, these loci become RdDM-dependent in *ddm1* somatic tissues, as indicated by the greatly lower CHH methylation levels in double mutants of DDM1 and genes involved in the canonical RdDM pathway, such as DRD1 (Fig 4B, *ddm1/drd1*, yellow bar) or RDR2 (Fig. 4B and C, *ddm1/rdr2*, red bars). In vegetative cells, embRdDM loci are co-targeted by both CMT2 and RdDM (Fig. 4D) since *drm* mutants and *cmt2* mutant vegetative cells show similar CHH methylation levels at embRdDM loci.

**Fig. 4.**
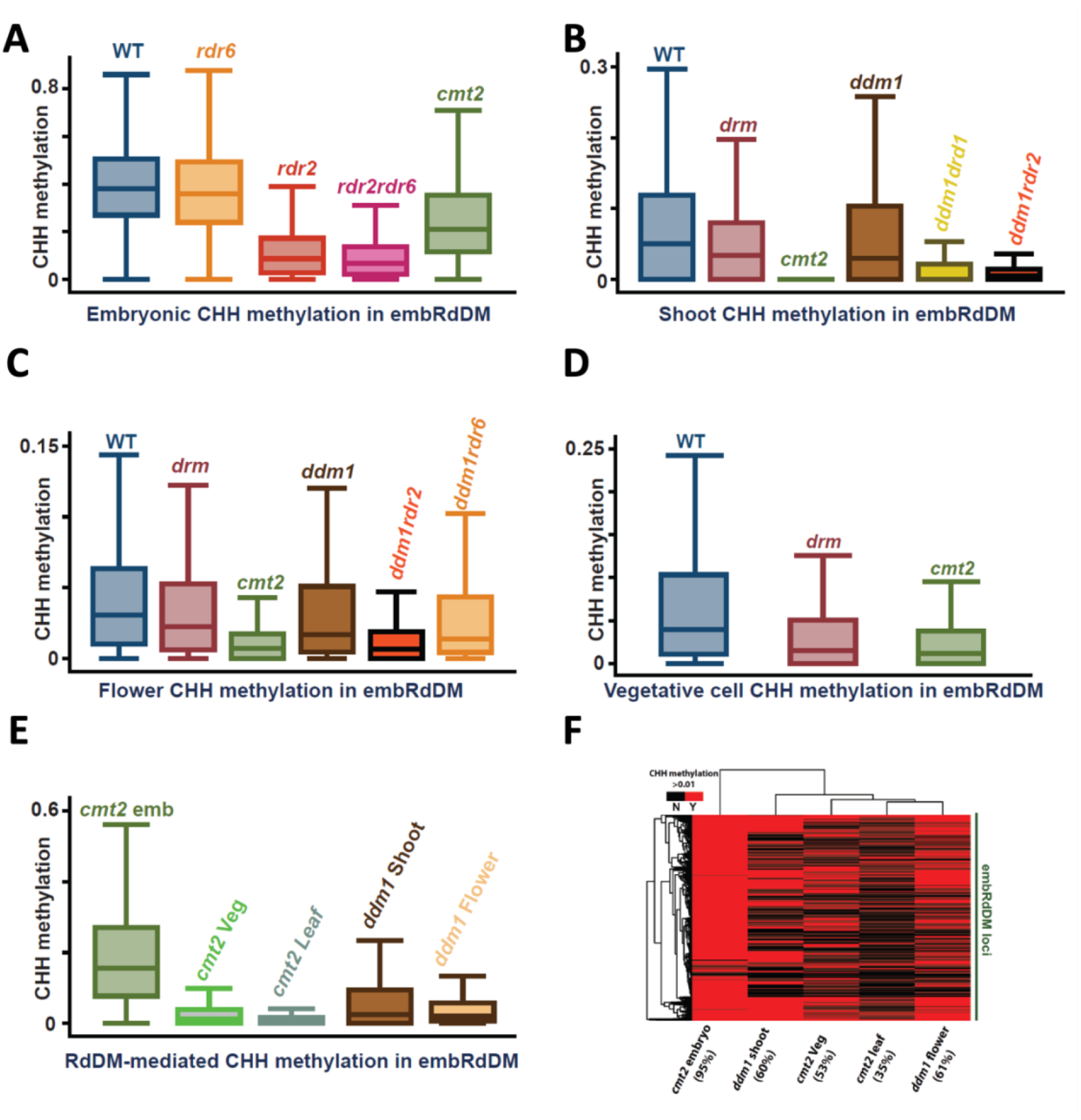
EmbRdDM loci are RDR2-dependent in embryos and partially overlap with RdDM targets in vegetative cells and *ddm1* somatic tissues. Box plots showing CHH methylation levels of 50-bp windows in embRdDM loci in WT, *cmt2*, *ddm1*, and RdDM-related mutants (*rdr2/6, drm, drd1*) in (A) mature embryos, (B) shoots, (C) flowers, and (D) vegetative cells. Only windows with at least 20 informative sequenced cytosines in each sample are included. (E) Box plots show RdDM-mediated CHH methylation levels in embRdDM loci in *cmt2* mature embryos, *cmt2* vegetative cells, *cmt2* leaves, *ddm1* shoots, and *ddm1* flowers. (F) Heatmap showing embRdDM loci possessing more than 1% RdDM-mediated CHH methylation in cmt2 mature embryos, cmt2 vegetative cells, cmt2 leaves, ddm1 shoots, and ddm1 flowers. Numbers show percentages of loci having CHH methylation greater than 0.01.

However, a significant portion of embRdDM loci are not catalyzed by RdDM in *ddm1* somatic tissues and vegetative cells. In *ddm1* shoots and flowers, where pericentric heterochromatin is more accessible (Zhong et al. 2021) ∼60% of embRdDM loci were methylated by RdDM machinery (i.e. CHH methylation > 0.01) (Fig. 4F). In *cmt2* mutant vegetative cells, where the effects of RdDM can be clearly observed: 53 % of CHH methylated loci (i.e. CHH methylation > 0.01) overlapped with embRdDM sites while the control *cmt2* mature embryos display 95% (Fig 4F). This is consistent with an earlier report showing that DDM1 is not expressed and pericentric heterochromatin is more accessible in the pollen vegetative cell (Borg et al. 2021; Slotkin et al. 2009). On the other hand, we observed that CHH methylation of pericentric-enriched vegetative-cell-specific accessible chromatin regions (VN-specific ACRs - (Borg et al. 2021)) is DRM-CMT2 co-dependent in leaves (Fig. S6A), and gradually becomes more DRM-dominant in vegetative cells and mature embryos (Fig. S6B and S6C). These findings suggest that VN-specific ACRs are likely more accessible for RdDM machinery in embryonic and vegetative cells than in leaf cells.

In addition to the canonical RDR2-dependent RdDM pathway, the transcription dependent RDR6-dependent RdDM pathway was reported to be involved in targeting derepressed full-length autonomous transposable elements in *ddm1* somatic tissues and vegetative cells (Martínez et al. 2016; Hsieh et al. 2016). To investigate whether embRdDM CHH hypermethylated loci are conferred similarly to those in vegetative cells and *ddm1* somatic tissues, we examined whether embRdDM loci were RDR6-dependent. We found that lack of RDR6 had only a marginal effect on the CHH methylation levels at embryonic novel RdDM targets, as well as overall at TEs (Fig. 4A), suggesting that the embRdDM loci in embryos are mostly RDR2-dependent, with the contribution of RDR6 being comparatively smaller. To assess whether sites of RDR6-mediated CHH methylation in *ddm1* somatic tissues and vegetative cells overlapped with embRdDM loci, we measured CHH methylation levels in embRdDM loci of *ddm1rdr6* flowers (publically available dataset accession GSM4305662) and found that additional *rdr6* mutation in *ddm1* flowers only slightly reduced the CHH methylation levels of embRdDM loci (Fig. 4C). Collectively, our results demonstrate that embRdDM loci are largely targeted by RDR2-dependent RdDM, which is different from *ddm1* somatic tissues and vegetative cells.

Our data show that the RDR2-dependent RdDM pathway targets embRdDM loci in *ddm1* somatic tissues and vegetative cells, however, the effect of RdDM in non-embryonic tissues is much weaker, since *cmt2* mature embryos showed much higher CHH methylation in these loci than *ddm1* somatic tissues and vegetative cells (Fig. 4E). Thus, although our analysis shows that embryos, vegetative cells and *ddm1* somatic tissues all display CHH hypermethylation and extended RdDM activities at common loci, around one half of embRdDM loci are unique to embryos, and the effect of RdDM is also much stronger in embryos (Fig. 4E and 4F). Since RDR6-dependent RdDM (i.e. transcription-dependent RdDM) is not the main mechanism for targeting embRdDM loci, it seems that the derepression of TEs is not the primary mechanism for extended RdDM activity in embryos. Instead, increased heterochromatin accessibility during early embryogenesis (Papareddy et al. 2020) may be the primary driver, which we explored next.

### Columella cells show CHH hypermethylation and enriched presence of small RNAs in new embryonic RdDM sites

The majority of Arabidopsis sporophytic tissues display similar CHH levels to leaves. However, columella root cap cells were reported to show a loss of heterochromatin and genome-wide CHH hypermethylation, accompanied by reduced transcriptional levels of heterochromatin-promoting genes (Kawakatsu et al. 2016). Columella cells, which are separated from the root apical meristem by only a few cell divisions, are thought to play a role in preventing harmful TE transcription by exporting small RNAs (sRNAs) to strengthen DNA methylation at meristem TEs, similar to the proposed function of the vegetative cell in pollen (Ibarra et al. 2012; Kawakatsu et al. 2016). Given these similarities between columella and embryo, we wondered whether the RdDM pathway could also target embRdDM loci in columella cells. To best capture the correlation between mature embryos and columella cells, we further merged 50-bp windows of the embRdDM loci within 300-bp regions, and defined regions at least 100-bp long and demonstrating a statistically significant difference in RdDM-mediated CHH methylation between leaves and embryos (Fisher’s exact test P-value < 10−5, and *cmt2* embryos have 10% more CHH methylation than *cmt2* leaves) at novel embryonic RdDM differentially methylated regions (embRdDM-DMRs).

The CHH methylation patterns of WT and *cmt2* embryos at embRdDM-DMRs were strikingly similar to columella root cap cells (Fig. 5A). In the same way that mature embryos were globally CHH hypermethylated compared to leaves (Fig. 2A and 5B, green and blue trace), columella cells exhibited global CHH hypermethylation compared to root stele cells, and both columella and mature embryos exhibited further enhanced CHH hypermethylation at embRdDM-DMRs. (Fig. 5B, brown and yellow trace). PCA analysis of CHH methylation profiles in embRdDM DMRs further revealed that columella and *cmt2* embryos exhibit similar DNA methylation profiles (Fig. 5C). These results are consistent with the lower expression of CMT2 as well as the absence of DDM1 protein in columella cells, which is required for CMT2’s function (Kawakatsu et al. 2016).

**Fig. 5.**
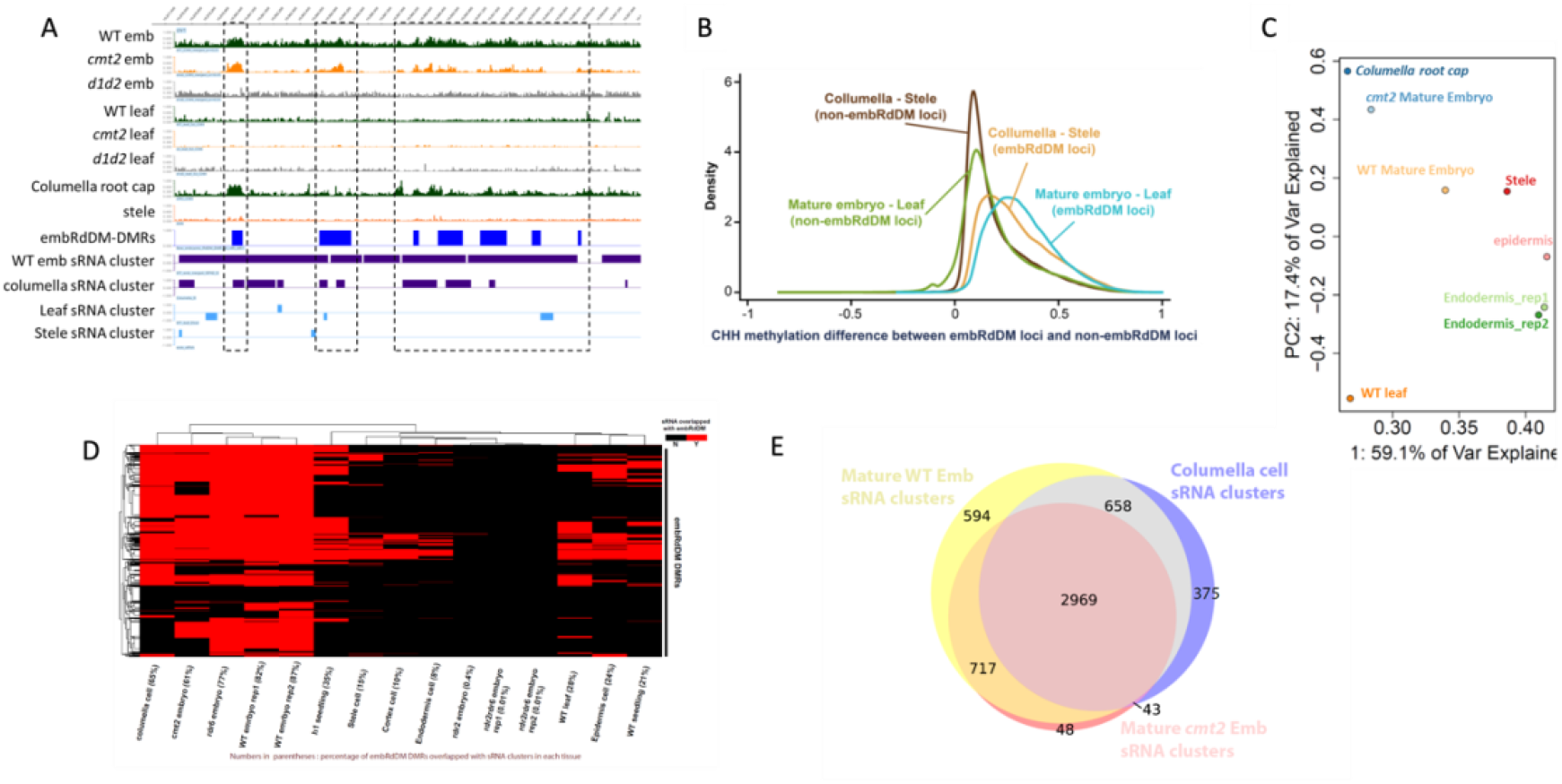
Columella cells also show CHH hypermethylation and enriched presence of small RNAs (sRNAs) in embRdDM DMRs. (A) Snapshots of CHH methylation in mature embryos and leaves of WT, *cmt2*, and *drm* mutants as well as WT columella and stele cells. embRdDM-DMRs are clustered from embRdDM loci as described in the main text (dark blue boxes). sRNA clusters generated by ShortStack are present in purple and light blue. Dashed-boxes indicate that WT and *cmt2* embryos show a similar CHH methylation pattern to columella cells in embRdDM DMRs. (B) Density plots showing the probability density of CHH DNA methylation differences between embryos and leaves, as well as between columella and stele cells in embRdDM DMRs and outside embRdDM DMRs. CHH methylation differences were calculated between 50-bp windows retaining at least 10% CHH methylation in either mature WT embryos or leaves in the embryo-leaf comparison. Only windows with more than 20 informative sequenced cytosines in both samples are included. The same cutoffs were used for the columella-stele comparison. (C) PCA analysis of CHH methylation profiles in embRdDM DMRs in different tissues. (D) Heatmap showing whether embRdDM DMRs are overlapped with sRNA clusters in different tissues. The red color indicates that embRdDM DMRs are overlapped with sRNA clusters by least 30% of the embRdDM DMR length. (E) Venn diagram showing the overlap of sRNA clusters among mature WT embryos, *cmt2* embryos, and columella cells in embRdDM DMRs.

In addition to the similarity of CHH methylation profiles between *cmt2* mature embryos and columella cells at embRdDM-DMRs, the presence of sRNAclusters in columella cells also correlated with embRdDM-DMRs (Fig. 5A). 65% of embRdDM-DMRs were overlapped with columella cell sRNA clusters, which is greatly enriched as compared to other root cell types, leaves and seedlings (Fig. 5D). Similarly, when examining WT embryo sRNA clusters at embRdDM loci, 74.6% (3686/4938) were overlapped with *cmt2* embryo sRNA clusters, and 73.4% (3627/4938) with columella cell sRNA clusters (Fig. 5E). Furthermore, we found that the sRNA abundance of embRdDM-DMRs is higher in the WT embryos and root columella cells than in other root cell types, seedlings, leaves and even pollen (within which the vegetative cell is likely the main contributor of sRNA) (Fig. S7A). With further analysis of the sRNA profiles of mature *rdr2*, *rdr6*, and *rdr2rdr6* embryos, we found that the production of small RNAs at embRdDM-DMRs is RDR2-dependent in embryos (Fig. 5D and Fig. S7A), as are the CHH methylation profiles of these mutants (Fig. 4A). Together, these results suggest that a conserved RDR2-dependent RdDM-mediated CHH methylation reconfiguration takes place in the chromatin-decondensed embryo and in columella cells.

H1 histone linker proteins are known to be enriched in heterochromatic regions and act to impede DNA accessibility for methyltransferases (Zemach et al. 2013). Loss of H1 is reported to be associated with CHH hypermethylation in heterochromatin, and H1 proteins are absent from CHH hypermethylated vegetative cells. We therefore hypothesized that embRdDM-DMRs could be associated with a lack of H1 proteins in embryos. To address this, we analyzed the *h1* root and h1 seedling methylomes but found the lack of H1 proteins did not further increase CHH methylation levels at embRdDM-DMRs (Fig. S7B and S7C). However, sRNA abundance in the embRdDM-DMRs was increased in *h1* seedlings (Fig. S7A), and the overlap of sRNAclusters with the embRdDM-DMRs also increased 14% in *h1* seedlings as compared to WT seedlings (Fig. 5D). These results indicate that the lack of H1 protein may only partially increase chromatin accessibility to RdDM machinery at embRdDM-DMRs, and is consistent with recent findings that heterochromatin remains largely intact and inaccessible to DNase I in *h1* seedlings (Choi et al. 2020). Also, since the embRdDM-DMR loci are more enriched in regions with intermediate H3K9me2 levels, the H1-enriched highly inaccessible heterochromatin is probably not the main target of embRdDM.

### embRdDM DMRs overlap with DEMETER targets and ovule siren loci

The DEMETER (DME) DNA glycosylase catalyzes genome-wide DNA demethylation in central and vegetative cells in plants (Hsieh et al. 2009; Ibarra et al. 2012; Schoft et al. 2011; Park et al. 2016) which is associated with sRNA production from small, euchromatic, AT-rich transposons, and at the boundaries of large transposons. sRNAs produced in the central and vegetative cells have been proposed to establish non-cell autonomous CHH methylation at corresponding DME-target loci in egg and sperm cells, respectively, via the RdDM pathway (Ibarra et al. 2012). Following double fertilization, the demethylated central cell genome is inherited by the endosperm and it has been suggested that maternal CG hypomethylated loci in endosperm generate sRNAs that promote embryo CHH methylation via RdDM (Hsieh et al. 2009; Mosher and Melnyk 2010)

To explore the possibility that endosperm may influence embryo DNA methylation, we compared embRdDM DMRs to DME targets in endosperm and vegetative cells to see if there was any overlap, finding that 50.4% (32516/64544) of embRdDM DMRs overlap with either endosperm DME DMRs or vegetative cell DME DMRs (Fig. 6A). Consistent with these results, CG hypermethylation of *dme* endosperm was more prevalent at embRdDM DMRs than non-embRdDM loci (Fig. 6B), mirrored by increased CG hypomethylation in wild type vegetative cell embRdDM loci as visualised by the positive shoulder on both density plots (Fig. 6C). Since both endosperm and vegetative cell DME targets overlap with embRdDM sites indicates that concordant chromatin states between DME targets and embRdDM sites is the reason for their shared locations. However, further work is needed to explore the question of whether gametophytic DME-mediated demethylation affects embryo CHH methylation.

**Fig. 6.**
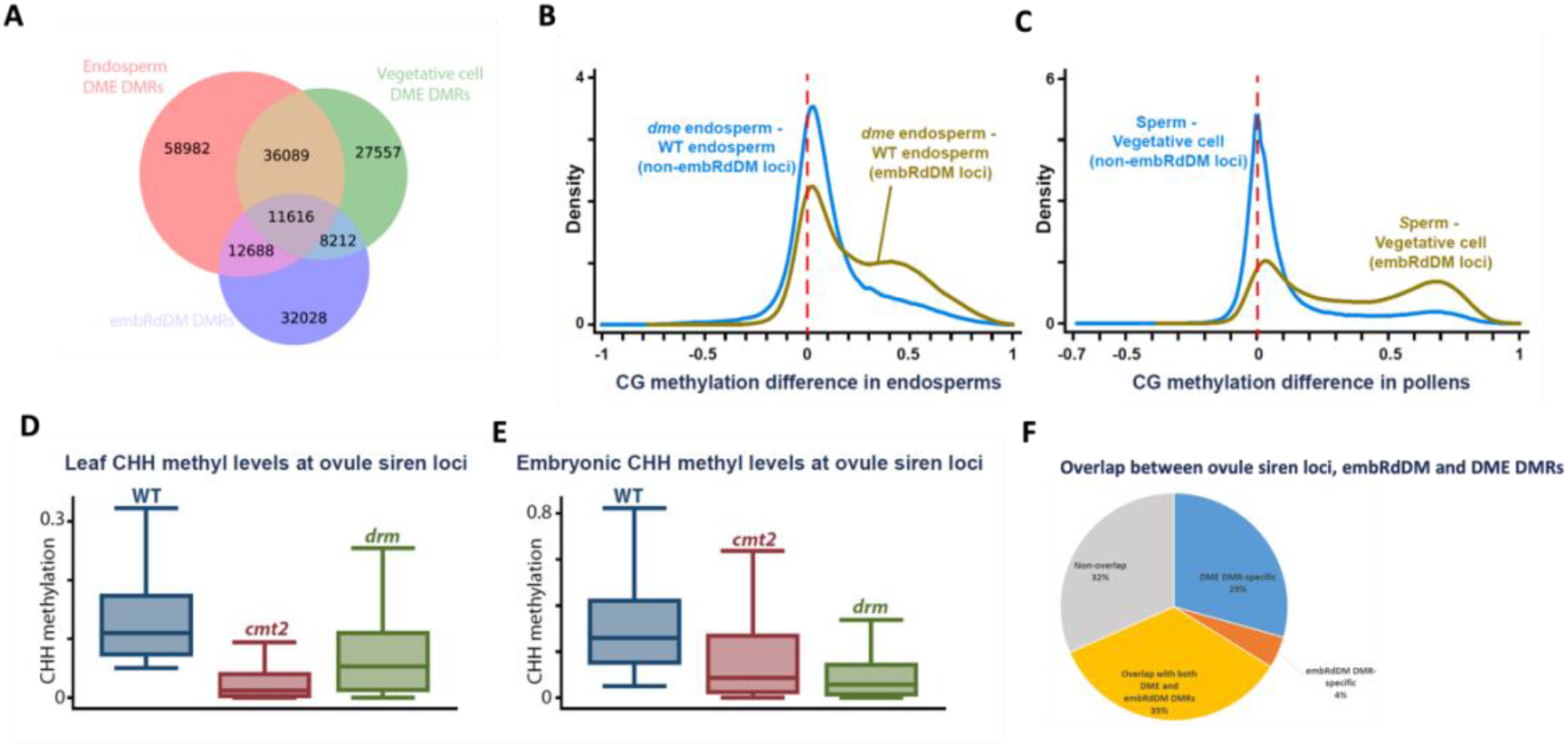
embRdDM DMRs, DEMETER targets, and ovule siren loci demonstrate striking overlap. (A) Venn diagram showing the overlap between endosperm DME DMRs, embRdDM DMRs, and vegetative cell DME DMRs. The overlaps were calculated based on 50-bp windows in each DMR. Density plots showing CG DNA methylation differences between (B) *dme* and WT endosperm, and between (C) sperm and vegetative cells within embRdDM DMRs and outside embRdDM DMRs. The CG methylation differences were calculated between 50-bp windows retaining at least 30% CG methylation in either WT or *dme* endosperm, or either sperm and vegetative cells in each comparison. Only windows with more than 10 informative sequenced cytosines in both samples are included. (D) Box plots show CHH methylation levels at ovule siren loci in leaves of WT, cmt2 and *drm* plants. (E) Box plots show CHH methylation levels at the ovule siren loci in embryos of WT, *cmt2* and *drm* plants. (F) Pie chart showing the overlap between ovule siren loci with different merged DME DMRs (endosperms and vegetative cells) and embRdDM DMRs.

Another potential contribution to embryonic CHH reprogramming may derive from epigenetic patterns in gametes that are inherited directly, or perhaps via non-cell autonomous signaling. Recent observations of CLSY-dependent reprogramming by RdDM in sperm and at siren loci in ovules are linked to high levels of tissue-specific siRNAs and CHH methylation (Long et al. 2021; Zhou et al. 2022). In leaves, CHH methylation at ovule siren loci is mainly regulated by CMT2 (Fig 6D). However, in embryos, there is a substantial skew towards CHH methylation dependence on RdDM, indicative of potential overlap between the epigenetic regulation of these loci between ovules and embryos (Fig. 6E). Consistent with this, we found that 39% of ovule siren loci overlap with embDMRs (Fig. 6F). Surprisingly, the merged DME DMRs of endosperm and vegetative cells show an even higher overlap with ovule siren loci (64% of siren loci are also DME targets, Fig. 6F). 35 % of siren loci were both DME targets and emRdDM DMRs. These data could either point towards gamete siRNA inheritance by the embryo, or a common chromatin conformation between these tissues that leads to similar epigenetic regulation.

## Discussion

In this paper we explore the mechanisms that lead to the programmed steady increase in CHH methylation throughout seed development. The early zygote has relatively low CHH methylation levels (Calarco et al. 2012; Ibarra et al. 2012; Papareddy et al. 2020), but CHH methylation in heterochromatic TEs gradually increases and surpasses that in leaves by the early torpedo stage of embryogenesis. The phenomenon of embryonic CHH build-up is an evolutionarily conserved epigenetic mechanism seen in Arabidopsis (Lin et al. 2017; Bouyer et al. 2017; Kawakatsu et al. 2017), *Brassica rapa* (Grover et al. 2020), soybean (Lin et al. 2017) and chickpea, (Rajkumar et al. 2020) and therefore may be a common feature of dicot seeds, potentially required to ensure TE silencing (Lin et al. 2017). Here, we identify both common and unique pathways of CHH methylation conferral between embryos and vegetative tissues. In particular, we observe that embryo-specific CHH methylation is mediated by RdDM, at loci overlapping with RdDM targets in the vegetative cell, *ddm1* mutants and the columella, that correlate with an increased capacity for RdDM in embryos, seemingly due to increased chromatin accessibility. Strikingly, we also observe a subset of embryonic RdDM sites that overlap with DME targets and ovule siren loci.

In general, our findings revealed that embryo CHH hypermethylation in Arabidopsis is predominately confined to the sequences already methylated in leaf tissues either by RdDM, CMT2 or by both machineries. However, by comparing the embryo methylomes of *cmt2*, *drm2*, *rdr2/6*, and wild type, we were able to delineate the timings of CHH hypermethylation activities between the various pathways. We found that CMT2-mediated CHH methylation begins during early embryogenesis, and RdDM-mediated CHH methylation predominates later, between early cotyledon and mature embryo, consistent with earlier reports (Bouyer et al. 2017; Papareddy et al. 2020). The activity of the RdDM pathway corresponds with the accumulation of siRNAs which peak at the mature embryo stage. These siRNAs are derived from short TEs, dispersed along pericentromeric and euchromatic DNA (Papareddy et al. 2020). The activity of CMT2 in embryogenesis is accompanied by a separate wave of siRNA accumulation, beginning just after fertilization and decreasing as seeds mature, derived from TEs that are generally longer and found in heterochromatic centromeres, and uniquely for embryos, derived from the entire TE sequence (Papareddy et al. 2020). It therefore seems likely that siRNA expression instigates both CMT2 and RdDM mediated CHH deposition, at distinct groups of TEs that respectively correspond to the canonical targets of CMT2 and RdDM. It is not clear why CHH deposition in early and late embryogenesis is mediated by CMT2 and RdDM, respectively, or why there is a switch during development.

Our data indicate that in large part, CHH hypermethylation in embryo occurs at sites that are also methylated in leaf and other somatic tissues, but at lower levels. We revealed a mechanistic switch in embryos, whereby many sites targeted by CMT2 in leaf became exclusively targeted or co-targeted by RdDM machinery. Since RdDM is known to target euchromatic TEs and edges of heterochromatic TEs due to chromatin accessibility, it seems likely that embryonic heterochromatin became accessible to the RdDM machinery via dynamic chromatin reprogramming. Indeed, Papareddy and colleagues showed that post-fertilization, zygotic chromocenters are decondensed, likely to facilitate rRNA production to ensure rapid embryo development (Papareddy et al. 2020). The initial low embryonic CHH methylation levels also imply that chromatin accessibility may be higher in embryos since DNA methylation is positively correlated with chromatin inaccessibility (Zhong et al. 2021; Noshay et al. 2021).

Furthermore, H1 depletion, which is linked to chromatin decondensation in the pollen vegetative cell to facilitate easiRNA production (He et al. 2019) and DEMETER demethylation in the central cell (Frost et al. 2018); also led to the induction of an embryo-like siRNA profile in leaves (Papareddy et al. 2020). The lack of H1 proteins did not further increase CHH methylation levels at embRdDM-DMRs, potentially because they were already saturated, however sRNA abundance at embRdDM-DMRs was increased, providing some supporting evidence that H1 configuration, and associated chromatin structure, in embryos contributes to their sRNA profile.

The high CHH methylation seen in the mature embryo resembles the high levels of CHH methylation seen in the pollen vegetative cell (Ibarra et al. 2012; Hsieh et al. 2016) and in the columella cells of the root tip (Kawakatsu et al. 2016), which may be linked to increased pericentric heterochromatin accessibility. One purpose for such decondensation might serve to facilitate rapid production of transcripts and ribosomal RNAs required to support the high metabolic needs of pollen tube elongation during germination (Mérai et al. 2014) and the rapid differentiation of columella during root growth (Gehring 2016). A similar biological purpose can be envisioned during the maturation stages of embryogenesis, when the embryo proper transitions from pattern formation to lipid and storage protein production, demanding high transcription and protein synthesis capability that persists through the mature embryo stage (Goldberg et al. 1994).

We focused on embryonic RdDM (‘embRdDM’) loci, which were specifically targeted by RdDM in embryo but not in leaf tissue. Many embRdDM loci overlapped with canonical DME target sites (Ibarra et al. 2012), as previously observed (Bouyer et al. 2017), which are also comprised of both euchromatic and heterochromatic TE targets, the latter being dependent on the chromatin remodeller FACT (Frost et al. 2018). Since the overlap was observed in both endosperm and vegetative cell DME targets, this suggests that similar chromatin conformation profiles between central and vegetative cells and embryo result in these overlapping targets between DME and embRdDM. We also observed a significant overlap between ovule siren siRNA loci and embRdDM sites, which could also be attributed to chromatin accessibility, potentially mediated by tissue-specific CLSY proteins, which regulate RdDM via locus-specific recruitment of Pol-IV (Zhou et al. 2018; Zhou et al. 2022).

Whether embryonic siRNAs were generated in situ by the embryo, or transported from endosperm or the surrounding seed coat has not been resolved, nor has their function fully elucidated (Bouyer et al. 2017; Grover et al. 2020; Kirkbride et al. 2019; Chakraborty et al. 2021). Of note, in *Brassica rapa*, maternal sporophytic RdDM, but not zygotic RdDM, is required for seed viability, showing that at least in this species, parental siRNA contributions are important for seed development, though maternal sporophyte RdDM was not sufficient to drive embryo hypermethylation (Chakraborty et al. 2021; Grover et al. 2020). The overlap between embRdDM loci and DME targets may reflect communication between embryo, endosperm and/or the seed coat, due to their intimate development within the seed. Endosperm also exhibits elevated CHH methylation, despite inheriting a globally CG-hypomethylated maternal genome (Hsieh et al. 2009; Ibarra et al. 2012). Indeed, abundant maternal-origin siRNAs have been reported from Arabidopsis siliques (Mosher et al. 2009) and whole seeds (Pignatta et al. 2014) and some maternally expressed imprinted genes are associated with siRNAs (Pignatta et al. 2014). However, 24 nt sRNAs generated by endosperm are distinct from those in embryos, being predominately genic in origin (Erdmann et al. 2017). Many siRNAs in Arabidopsis endosperm are produced from the maternal genome and have corresponding matching seed coat siRNAs (Kirkbride et al. 2019). Thus, it remains a possibility, at least in Arabidopsis seeds, that some sporophytic siRNAs are transported into developing filial tissues to influence their DNA methylation and transcription, as a means to reinforce TE silencing, or perhaps to manifest maternal influence over the zygote.

## Materials and Methods

### Isolation of Arabidopsis embryos

Siliques at approximately 7-12 DAP were chosen and dissected and seeds mounted on sticky tape. Mounted seeds were immersed in dissection solution [filter-sterilized 0.3 M sorbitol and 5 mM (pH 5.7) MES] and dissected by hand under a stereomicroscope using fine forceps (Inox Dumont no. 5; Fine Science Tools) and insect mounting pins. The seed coat was discarded, and debris was removed by washing collected embryos six times with dissection solution under the microscope.

### Bisulfite sequencing library preparation

Genomic DNA samples of embryos were extracted by the Qiagen DNeasy Plant Kit following the manufacturer’s protocol. Single-end bisulfite sequencing libraries for Illumina sequencing Library preparation was performed as described in (Frost et al. 2018). In brief, 50 ng of genomic DNA was fragmented by sonication, end repaired and ligated to custom-synthesized methylated adapters (Eurofins MWG Operon) according to the manufacturer’s instructions for gDNA library construction (Illumina). Adaptor-ligated libraries were subjected to two successive treatments of sodium bisulfite conversion using the EpiTect Bisulfite kit (Qiagen) as outlined in the manufacturer’s instructions. The bisulfite-converted library was split between two 50 ul reactions and PCR amplified using the following conditions: 2.5 U of ExTaq DNA polymerase (Takara Bio), 5 μl of 10X Extaq reaction buffer, 25 μM dNTPs, 1 μl Primer 1.1 and 1 μl multiplexed indexing primer. PCR reactions were carried out as follows: 95°C for 3 minutes, then 14-16 cycles of 95 °C 30 s, 65 °C 30 s and 72 °C 60 s. Enriched libraries were purified twice with AMPure beads (Beckman Coulter) prior to quantification with the Qubit fluorometer (Thermo Scientific) and quality assessment using the DNA Bioanalyzer high sensitivity DNA assay (Agilent). Sequencing on either the Illumina HiSeq 2000/2500 or HiSeq 4000 platforms was performed at the Vincent J. Coates Genomic Sequencing Laboratory at UC Berkeley.

### Methylome data processing

Bisulfite-seq paired-end reads were mapped to the TAIR10 reference genome by Bismark (v.0.17.0) with the command: --bowtie1 --un --multicore 6 --chunkmbs 6400 -q -n 2 -l 100. Methylation analysis was performed by bismark_methylation_extractor with the command: -p --no_overlap --bedGraph --counts --buffer_size 10G --cytosine_report –CX -split_by_chromosome.

### Methylome data analysis

The CHH methylation information from two biological replicates of WT, *cmt2*, *drm1drm2*, *rdr2*, *rdr6* and *rdr2rdr6* mature embryos were merged after Fig.1 when the CHH levels between two replicates were not significantly different in a single-base resolution (Fischer exact test p-value>0.01). CHH methylation machinery classification was determined by the following steps: (1) WT CHH methylation must be greater than 0.01 in 50-bp windows. 50-bp windows where the leaf CHH methylation is less than 0.01 were defined as unmethylated loci (2) only 50-bp windows where WT, *cm2*, *drm* leaves and embyos all have at least 20 informative reads were kept (3) defined the cmt2-CHH-index = (WT CHH – *cmt2* CHH)/(WT CHH – *cmt2*)+ (WT CHH – *drm* CHH) for each 50bp window. When *cmt2* or *drm* mutants have more residual CHH methylation than WT, the differences between mutants and WT were set to zero. (4) DRM-dominant loci were defined when cmt2-CHH-index<=0.4, CHH methylation levels are significantly different between WT and *drm* mutants (Fischer exact test p-value<0.01), and *drm* mutants lost more than 50% of WT CHH methylation (5) CMT2-dominant loci were defined when cmt2-CHH-index>=0.6, CHH methylation levels are significantly different between WT and *cmt2* mutants (Fischer exact test p-value<0.01), and *cmt2* mutants lost more than 50% of WT CHH methylation (6) DRM-CMT2 co-targeting loci were defined when 0.6>cmt2-CHH-index>0.4, CHH methylation levels are significantly different between WT and *drm* mutants as well as WT and *cmt2* mutants (Fischer exact test p-value<0.01 for both comparisons). The classification was performed on both the leaf and the mature embryo datasets. The new embryonic RdDM (embRdDM) loci are the sum of: 1. 50-bp windows that are unmethylated in leaves but become DRM-dominant targets or DRM-CMT2 co-targeting targets in embryos; 2. 50-bp windows that are CMT2-dominant in leaves but become DRM-dominant targets or DRM-CMT2 co-targeting targets in embryos.

### Small RNA sequencing library preparation

We extracted total RNA from mature embryos of WT, *rdr2*, *rdr6*, *rdr2rdr6*, *cmt2* and *rdr6* mutants with Trizol (Invitrogen, cat. no. 15596026). Total RNA samples were treated with the TURBO DNA-free Kit to remove contaminating genomic DNA. RNA was then subjected to sRNA library preparation according to the manufacturer’s protocol (Illumina, cat. no. RS-200-0012 and RS-200-0024).

### Small RNA sequencing data processing

Adapter sequences were trimmed from the reads and resultant trimmed reads were sorted into size classes from 18 nt to 28 nt using custom Python scripts (Rodrigues et al. 2021). The trimmed reads were mapped to TAIR10 pre-tRNA and rRNA sequences by BOWTIE and the unmapped reads were used for the further analysis (option: -v 0 -B 1 --best --un). The non-tRNA and rRNA reads were mapped to the TAIR10 reference genome and the small RNA clusters were called by ShortStack (option: --mincov 2.0rpm --nohp --bowtie_cores 1 --mismatches 0).

### Published Genomic Datasets utilised

DNA methylation data for WT, *cmt3*, *cmt2*, *drm1drm2 (drm*), and ChIP-seq data for H3K9me2 and histone H3 in leaf tissues are from Stroud et al. 2014. DNA methylation data for WT, *cmt2* and *drm* vegetative cells and WT sperm cells are from Hsieh et al., 2016. Methylome data for root columella, stele, epidermis, endodermis, and cortex cells are from Kawakatsu et al., 2016. The small RNA datasets from columella, stele, epidermis, endodermis, and cortex cells are from (Breakfield et al. 2012). DNA methylation and small RNA datasets for WT and h1 seedlings are from (Choi et al. 2021; Choi et al. 2020). DNA methylation data for WT and *h1* root are from (Zemach et al. 2013). Embryo H3K9me2 and histone H3 ChIP-seq data are from (Parent et al. 2021). DNA methylation data for WT, *cmt2*, *drm2*, *ddm1*, *ddm1rdr2*, *ddm1rdr6*, and *ddm1drd1* flower and shoot tissues are from (Panda et al. 2016; Zemach et al. 2013). DNA methylation data for generating DME-WT endosperm DMRs are from (Hsieh et al. 2009). DNA methylation data for generating sperm-vegetative-cells DMRs are from (Ibarra et al. 2012).

### Data availability

Sequencing data will be deposited into NCBI’s GEO. Accession TBC.

## Acknowledgements

This work was supported by NSF Grant IOS-1025890 (to R.L.F. and D.Z.), NIH Grant GM69415 (to R.L.F. and D.Z.), a faculty scholar grant from HHMI and the Simons Foundation 55108592 (to D.Z.), NIFA Hatch Project 02413 (to T.-F.H.) and NSF Grant MCB-1715115 (to T.-F.H.), and NRF of Korea Grants 2020R1A2C2009382 and 2021R1A5A1032428 (to Y.C.). We thank Christina Wistrom and formerly Barbara Rotz for their management of the UC Berkeley Oxford Tract greenhouse facility, and we are grateful to Christian Ibarra for his advice on developing seed dissection. This work used the Vincent J. Coates Genomics Sequencing Laboratory at UC Berkeley, supported by NIH Instrumentation Grant S10 OD018174.

---

**Fig. S1.**
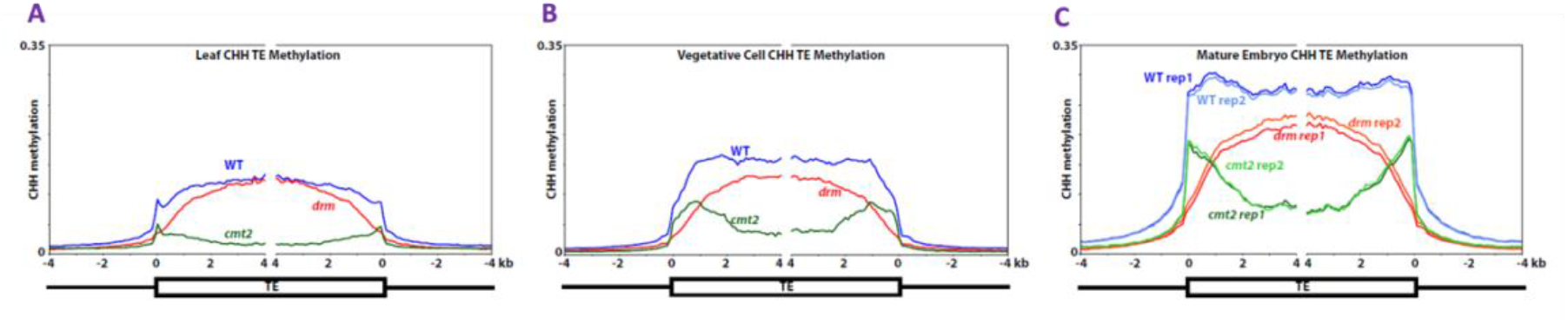
CHH hypermethylation is maintained similarly in TEs in leaves, vegetative cells and mature embryos. Ends analyses showing methylation within each 100-bp interval averaged and plotted from 4 kb away from the annotated TE (negative numbers) to 4 kb into the annotated region (positive numbers). CHH methylation in (A) Leaf TEs, (B) Vegetative Cell TEs and (C) Mature embryo TEs in WT, *cmt2* and *drm* mutants.

**Fig. S2.**
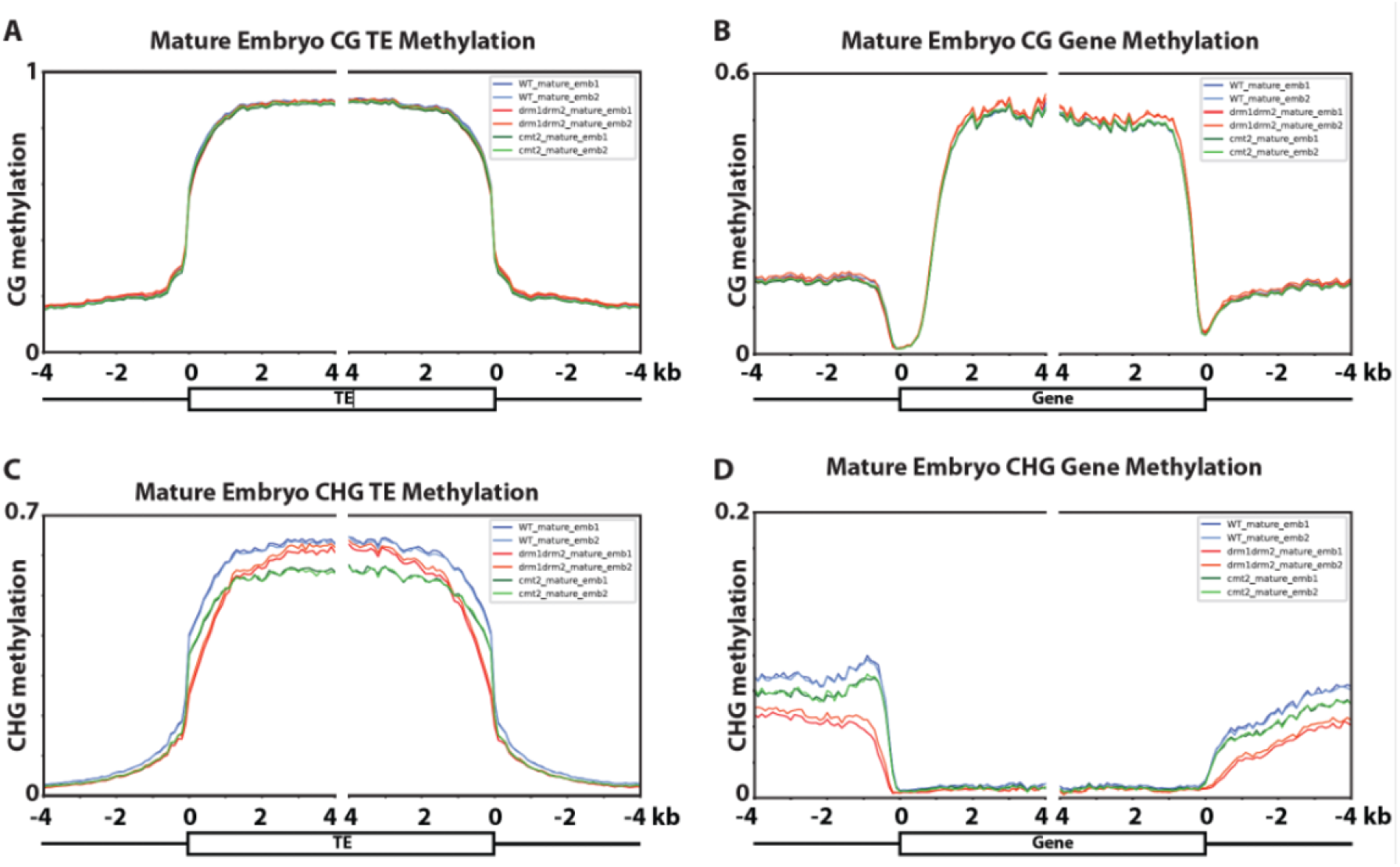
Embryonic CG and CHG methylation at genes and TEs in mature embryos. Ends analyses showing methylation within each 100-bp interval averaged and plotted from 4 kb away from the annotated TE (negative numbers) to 4 kb into the annotated region (positive numbers). (A) TE methylation in mature embyros is unchanged in *drm* and *cmt2* mutants (B) Gene body methylation is slightly elevated in *drm* mutant embryos (C) Loss of CMT2 and DRM resulted in a slight decrease of embryonic CHG methylation in TEs, but not in gene bodies (D).

**Fig. S3.**
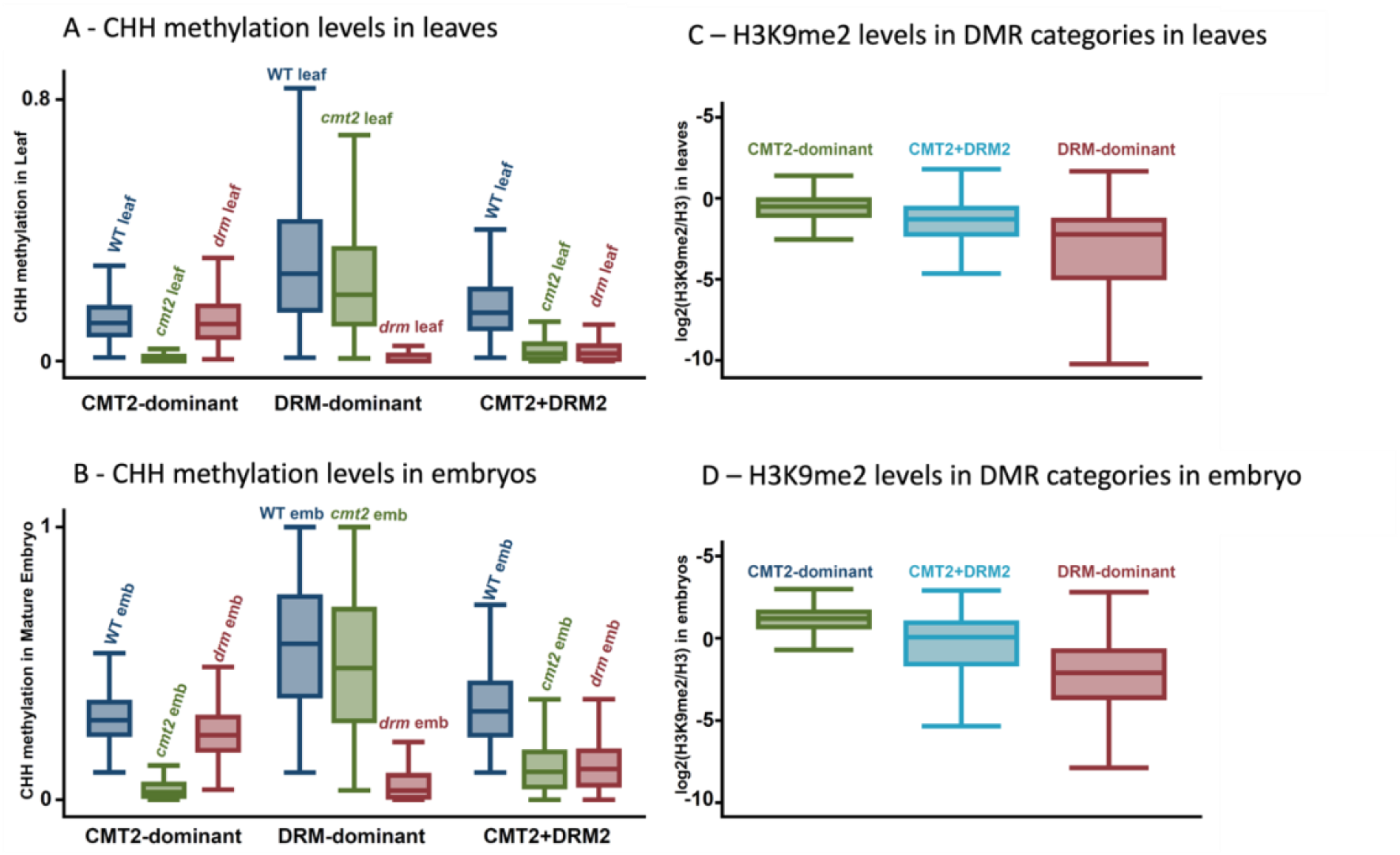
Validation of CHH DMRs in leaves and embryos, and the H3K9me2 levels of these DMRs. (A) Box plot showing CHH methylation levels in leaves and (B) in embryos for each mutant genotype, demonstrating the validation of respective methylation DMRs (C) H3K9me2 levels in leaves and (D) in embryos (D) at CMT2-dominant DMRs, CMT2+DRM2 shared DMRs and DRM dominant DMRs.

**Fig. S4.**
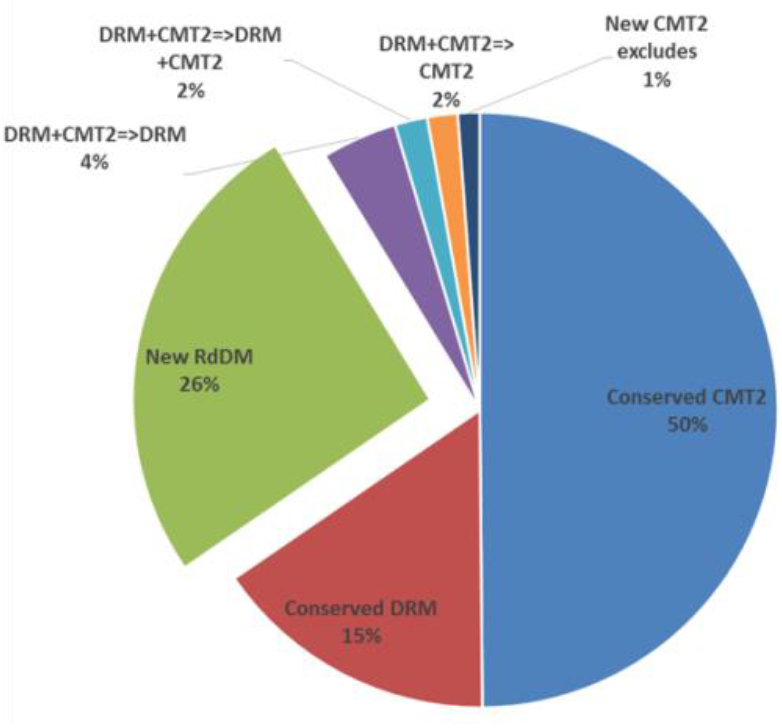
Summary of mechanisms mediating embryonic CHH hypermethylation loci in mature embryos. The majority (50 %) of CHH hypermethylated sites in embryos are conserved with those in leaves, albeit exhibiting lower methylation levels in leaves, and are conferred by CMT2. The next largest group (25 %) are new (or undetectable in leaves) CHH methylated sites in embryos, conferred by RdDM. The remainder are conserved DRM (RdDM) sites shared in embryos and leaves (15 %) and other smaller groups representing combinations of the above.

**Fig. S5.**
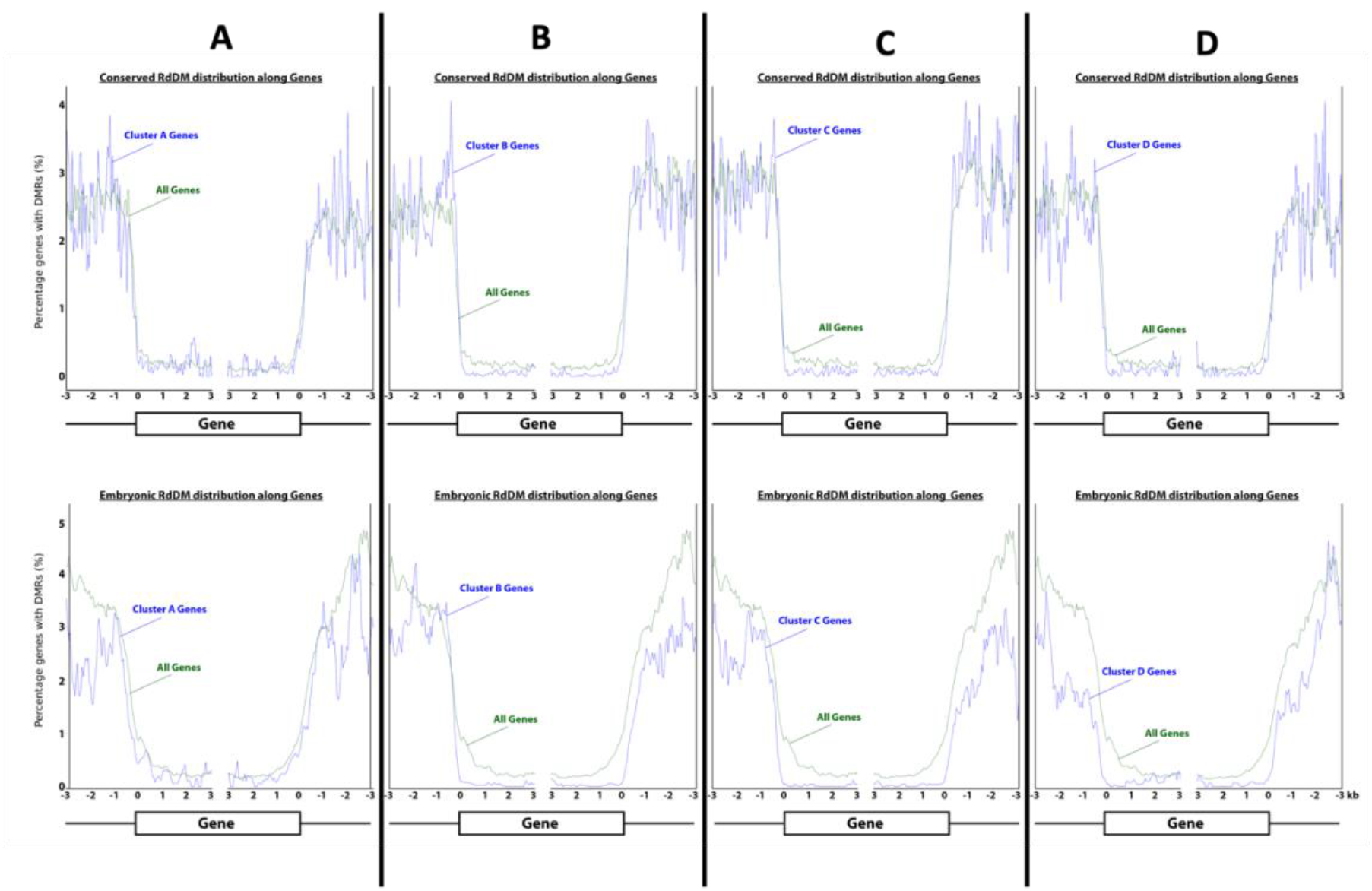
Neither conserved nor embryonic RdDM are associated with gene expression patterns. Distribution of embRdDM loci and consvRdDM loci along genes with different expression patterns during embryogenesis – (A) gene cluster with highest expression during the pre-cotyledon phase; (B): gene cluster for which expression decreases between transition and mature phases; (C): gene cluster with highest expression during the transition phase; (D): gene cluster with highest expression during the mature phase

**Fig. S6.**
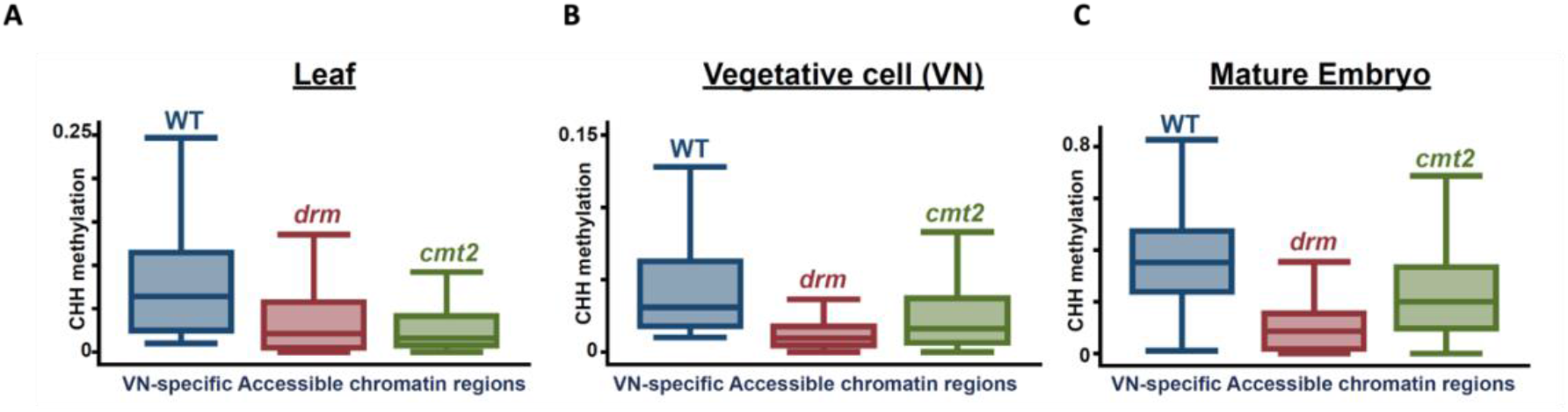
VN-specific ACRs are likely more accessible for RdDM machinery in embryonic and vegetative cells than in leaf cells. Box plots showing CHH methylation levels of Wildtype (WT), *drm* and *cmt2* mutants at vegetative cell specific accessible chromatin regions in (A) leaf, (B) vegetative cell, and (C) mature embryo in VN-specific accessible chromatin regions (derived from Borg at al., 2021)

**Fig. S7.**
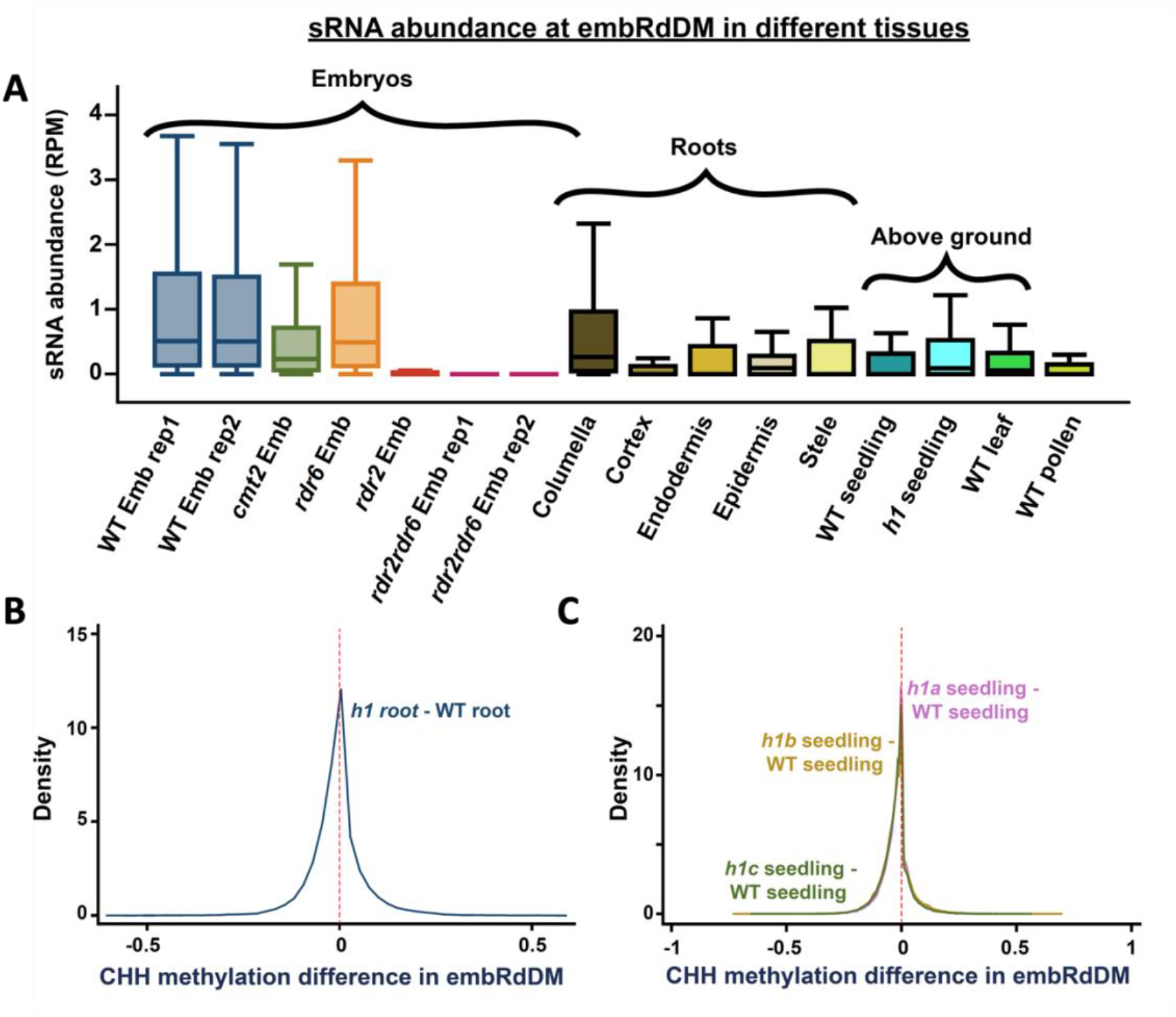
sRNA abundance of embRdDM-DMRs is higher in the WT embryos and root columella cells than in other root cell types, seedlings, leaves and pollen; lack of H1 protein marginally increases chromatin accessibility to RdDM machinery at embRdDM-DMRs. (A) Box plots show sRNA abundance at embRdDM DMRs in different tissues of WT, RdDM, and *cmt2* mutant plants. Kernel density plots of CHH methylation differences at embRdDM DMRs (B) between *h1* triple mutant and WT roots, and (C) between *h1a/b/c* and WT seedlings. CHH methylation differences were calculated between 50-bp windows with at least 20 informative sequenced cytosines in both samples.

**Table S1.** Summary of embryonic CHH hypermethylated loci according to their targeting mechanisms in leaves and embryos. Shaded rows indicate conserved DRM sites (first shaded row) and conserved CMT2 sites (second shaded row). There are high levels of conservation of CHH methylation mechanisms between leaves and embryos.

| Emb CHH - Leaf CHH $\geq 0.1$ | | | | |
| --- | --- | --- | --- | --- |
| Leaf (% in leaf categories ) |  | Emb categories | % based on leaf categories | # of 50bp windows |
| 11% | Lowly methylated | DRM | 59.4 | 10963 |
|  | Lowly methylated | CMT2 | 6.8 | 1250 |
|  | Lowly methylated | DRM+CMT2 | 33.8 | 6239 |
| 16% | DRM | DRM | 97.1 | 26584 |
|  | DRM | CMT2 | 0.3 | 92 |
|  | DRM | DRM+CMT2 | 2.5 | 688 |
| 66% | CMT2 | DRM | 7.0 | 7798 |
|  | CMT2 | CMT2 | 76.0 | 85182 |
|  | CMT2 | DRM+CMT2 | 17.1 | 19168 |
| 7% | DRM+CMT2 | DRM | 54.5 | 7001 |
|  | DRM+CMT2 | CMT2 | 21.9 | 2818 |
|  | DRM+CMT2 | DRM+CMT2 | 23.5 | 3023 |

